# Epidermal cell fusion promotes the transition from an embryonic to a larval transcriptome in *C. elegans*

**DOI:** 10.1101/2024.05.22.595354

**Authors:** Owen H. Funk, Daniel L. Levy, David S. Fay

**Author notes:** Corresponding authors: David S. Fay, University of Wyoming, Department of Molecular Biology, 1000 E. University Avenue, Laramie, WY 82071, Fay Daniel L. Levy, University of Wyoming, Department of Molecular Biology, 1000 E. University Avenue, Laramie, WY 82071.

## Abstract

Cell fusion is a fundamental process in the development of many multicellular organisms but its precise role in gene regulation and differentiation remains largely unknown. The *Caenorhabditis elegans* epidermis, which is comprised of multiple syncytial cells in the adult, represents a powerful model for studying cell fusion in the context of animal development. The largest of these epidermal syncytia, hyp7, integrates 139 individual nuclei through processive cell fusion mediated by the fusogenic protein EFF-1. To explore the role of cell fusion on developmental progression and associated gene expression changes, we conducted transcriptomic analyses of *eff-1* fusion-defective *C. elegans* mutants. Our RNAseq findings showed widespread transcriptomic changes including the enrichment of epidermal genes and molecular pathways involved in epidermal function during development. Single-molecule fluorescence *in situ* hybridization further validated the observed altered expression of mRNA transcripts. Moreover, bioinformatic analysis suggests that fusion may play a key role in promoting developmental progression within the epidermis. Our results underscore the significance of cell–cell fusion in shaping transcriptional programs during development.

**Summary Statement:** *eff-1* meditated cell fusion drives transcriptomic progression from an embryonic to larval state in the *C. elegans* epidermis, without which developmental progression is delayed.

## Introduction

Multinucleate syncytial cells are evolutionarily conserved and are found throughout the tree of life in physiologically healthy tissues and in disease states such as cancer (1-6). The need for a syncytium varies among organisms and tissues; multiple nuclei may be required to metabolically support larger cells or to facilitate the rapid production of large amounts of mRNA for developmental processes or in response to extracellular cues (7-11). There are multiple paths by which multinucleate syncytia arise, including cell–cell fusion of multiple mononucleate cells (1, 12-15). Cellular fusion necessarily merges the cytoplasm of two distinct cells, but how mixing of cytoplasmic factors changes the regulatory environment of individual nuclei and thus affects the transcriptome within a newly formed syncytium is poorly understood (16-19).

The *C. elegans* epidermis (also termed hypodermis) is an excellent model to address questions related to the function of syncytial nuclei. *C. elegans* has a well-characterized and largely invariant cell lineage in which over a third of somatic nuclei ultimately reside within syncytial cells (14, 20). The adult worm epidermis is composed of multiple syncytial cells, the largest of which is hyp7, which surrounds most of the adult body (14). hyp7 forms through progressive rounds of cell fusion, beginning with 23 embryonic fusion events that create the initial larval hyp7 prior to hatching. This process is completely dependent on cell–cell fusion mediated by the fusogen EFF-1 and is followed by the addition of 116 additional nuclei through EFF-1–mediated fusion events during larval development (21-23).

*eff-1* was initially discovered through genetic screens as the lone factor that is both necessary and sufficient to induce cell fusion in the *C. elegans* epidermis (22, 23). Mutations in *eff-1* lead to the disruption of embryonic fusion events in the developing hypodermal syncytium, leaving the hypodermal progenitors unfused and embryonic cell boundaries intact. *eff-1* mutant worms exhibit variable morphological and developmental defects, highlighting the importance of cell fusion for proper body morphogenesis (21, 23-25).

Here we explore the role of embryonic cell fusion in gene expression and differentiation in the developing epidermal syncytium through RNAseq of worms containing reduction-of-function mutations in EFF-1, which block most epidermal fusion events in the embryo. Reduced fusion led to an increase in embryo-associated gene transcripts concurrent with a decreased expression of larval-associated transcripts in genes expressed specifically in the epidermis. These findings suggest a developmental delay in the acquisition of epidermal cell fates and a possible role for cell fusion in activating gene regulatory programs associated with epidermal differentiation, adding to the growing list of functions for cell fusion and syncytialization.

## Results and Discussion

### *eff-1*–mediated cell fusion is essential for normal developmental progression and morphology in *C. elegans*

The discovery of EFF-1 as the primary fusogen governing most epidermal cell fusions in *C. elegans* originated from classical mutagenesis screens (21-23). This included two mutations—P183L (*hy21*) and S441L (*oj55*)—that exhibit variable penetrance fusion defects and temperature sensitivity. Based on available data, both mutations appear to be partial reduction-of-function mutations rather than complete knockouts and would be expected to contain background mutations as a result of the mutagenesis, as well as additional differences in strain backgrounds (23). We therefore generated strains containing an *eff-1* putative null mutation using CRISPR (P37Stop), which places two stop codons in exon 1 that is present in all *eff-1* isoforms (**Fig. 1A**). As discussed below, this precise modification may lead to a stronger reduction of function of *eff-1* as compared with the historical alleles and allows for a cleaner comparison to our wild type (WT; N2) fusion-competent control.

**Figure 1.**
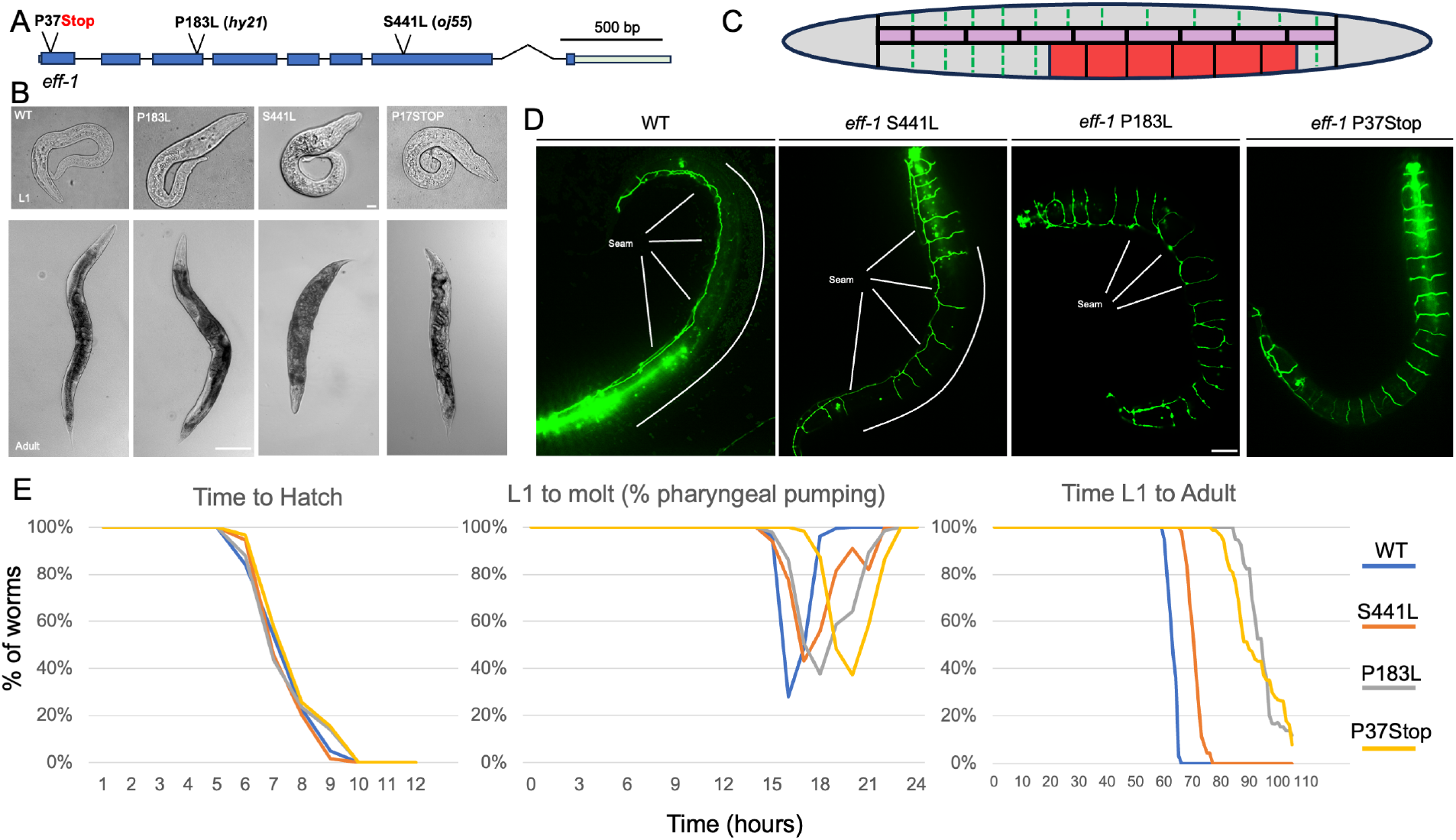
*eff-1* mutant worms display morphological defects and developmental delay. A) *eff-1* gene locus with CRISPR edit P37Stop and mutant alleles P183L (*hy21*) and S441L (*oj55*) indicated. B) Brightfield imaging of L1 and adult N2 and *eff-1* mutants as indicated. Scale bars: L1, 10µm; adult, 100µm. C) Schematic of L1 hypodermis with *eff-1*–mediated fusion events marked with dashed green lines, seam-cells in pink and P cells in red. D) Representative images of AJM-1::GFP fluorescence, a marker of apical cell–cell junctions in the epidermis, in N2 (WT) and *eff-1* mutant L1 larvae; lateral seam cells are labeled. E) Quantification of developmental timepoints in N2 and *eff-1* mutants. Time to first molt was measured via the cessation of pharyngeal pumping. Time to hatch and adulthood was measured by locomotion and sexual maturity, respectively. n of > 40 per condition and timepoint.

Consistent with previous reports, *eff-1*(P37Stop) mutants displayed characteristic morphological defects, including a bulging anterior region, tail spike abnormalities, and a dumpy (Dpy) phenotype that persisted into adulthood (**Fig. 1B**) (23). To visualize fusion defects in *eff-1* mutants we imaged apical junctions using AJM-1::GFP and confirmed that *eff-1(*P37Stop) mutants have an unfused epidermis that persists into adulthood (**Fig. 1C,D**). We note that the original characterization of the *eff-1* mutants reported a complete lack of fusion during late embryonic stages but with partial fusion occurring during larval development at temperatures below 25°C (23).

We next characterized the developmental timing of *eff-1*(P37Stop) and the historical mutants from hatching to first molt and to reproductive adulthood. Like *eff-1*(P183L) and *eff-1*(S441L), *eff-1*(P37Stop) exhibited developmental delays in the time to first molt and the time to reach adulthood, although the delays in *eff-1*(P183L) and *eff-1*(P37Stop) appeared more pronounced than *eff-1*(S441L) (**Fig. 1E**). Additionally, all three *eff-1* mutants were less synchronous than WT and we observed a low frequency of larval arrest in the *eff-1*(P183L) and *eff-1*(P37Stop) mutants.

### Disruption of cell fusion in embryonic development leads to consistent, widespread transcriptomic changes in *eff-1* mutants

The transition from embryo to larva is accompanied by widespread transcriptional rewiring as the newly hatched L1s begin locomotion and feeding and as many tissues and cell types acquire their terminal post-mitotic fate (26-29). To examine the role of cell fusion in the embryonic-to-larval transition, we conducted RNAseq on synchronized early L1-stage WT, *eff-1*(P37Stop), *eff-1*(S441L), and *eff-1*(P183L) L1 animals. For *eff-1*(P37Stop), differential expression analysis revealed 2046 upregulated and 912 downregulated differentially expressed genes (DEGs) relative to WT with a false-discovery rate (FDR) cutoff of <0.05 and log_2_(fold change) cutoff of >0.5 or <-0.5 (**Fig. 2A, File S1**). Additionally, we found 3534 upregulated and 2558 downregulated DEGs in *eff-1*(S441L) mutants, and 6499 up and 6829 down DEGs in *eff-1*(P183L) mutants (**Fig. S1A, File S1**). The larger number of DEGs observed in the previously isolated mutant strains is not entirely unexpected given that these strains were generated by random mutagenesis and that the parental strains used to generate these alleles may differ somewhat from our N2 reference strain. As such, some of the transcriptomic changes in these strains are likely attributable to background variations.

**Figure 2.**
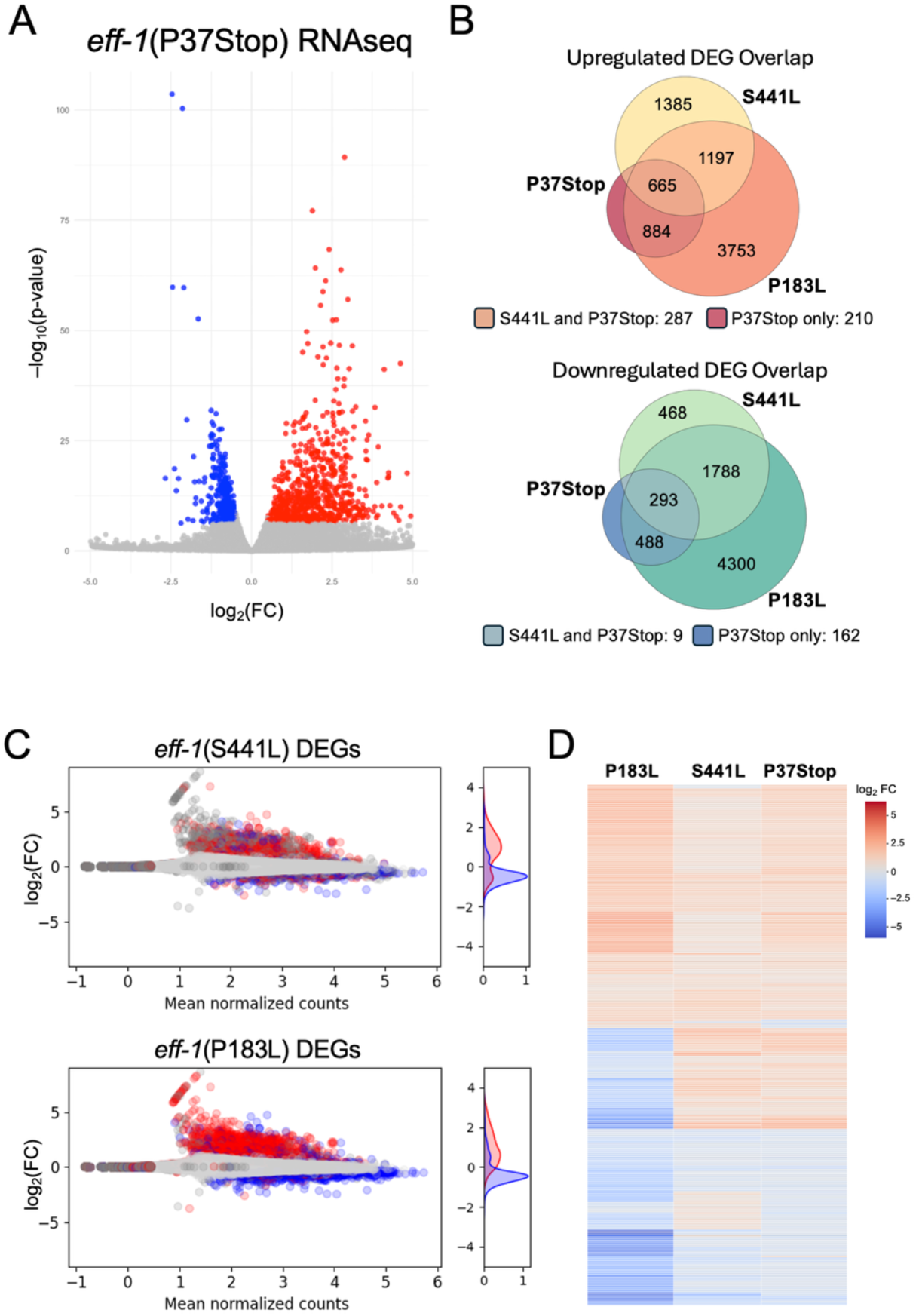
RNAseq of multiple *eff-1* mutants indicates a common transcriptome response to fusion disruption. A) Volcano plot of top DEGs (downregulated in blue, upregulated in red) for *eff-1*(P37Stop). B) Overlap of significant (adjusted p-value < .05, log_2_FC < -0.5 or > 0.5) DEGs from P37Stop, S441L, and P183L. C) MA (minus average; Bland-Altman) plot of *eff-1*(P37Stop) mutants with upregulated (red) and downregulated (blue) DEGs from S441L and P183L relative to WT. DEGs up- and downregulated in one mutant are more often found differentially expressed in the same direction in the other mutant. Right-side inset shows density of colored (shared DEGs) points D) Heatmap of log_2_-transformed fold changes (FCs) of top 200 DEGs shared between the *eff-1* mutants.

Despite potential background differences, we observed a strong overlap between *eff-1*(P37Stop), *eff-1*(S441L), and *eff-1*(P183L), with 958 shared DEGs that were differentially regulated in the same direction (293 downregulated and 665 upregulated) (**Fig. 2B,C**). Furthermore, log_2_-transformed fold-change data for the top DEGs from all three mutant strains also showed high concordance, with top genes shifting in the same direction regardless of the *eff-1* mutation (**Fig. 2D**). Euclidian distances between samples also confirmed transcriptome-wide similarities between *eff-1* point mutations and P37Stop as compared with WT (**Fig. S1B**). Notably, the P183L transcriptome appears highly divergent, even more so than the P37Stop despite being only a point mutation. We suspect this is due to additional background mutations introduced in the original derivation of this strain, further highlighting the utility of a clean, engineered knockout.

Taken together, these data indicate a shared core transcriptomic response to the absence of epidermal cell fusion in L1 larvae, regardless of the specific *eff-1* allele or genetic background. The substantial overlap in both the identity of DEGs and the directionality of their fold changes across all *eff-1* mutant conditions suggests a conserved transcriptional signature linked directly to the disruption of epidermal syncytialization. This highlights a fundamental set of gene expression changes that is consistently triggered by the failure of cells to fuse in the developing epidermis.

### Disruption of *eff-1*–mediated fusion alters epidermal cell expression

The hyp7 syncytium has many functions including the production of a new cuticle at each molting cycle, which occurs via the synthesis and secretion of apical extracellular matrix (aECM) proteins that include many collagens (30-34). Consistent with a disruption of these functions, Gene Ontology (GO) enrichment analysis showed a substantial number of terms associated with epidermal-specific processes, specifically extracellular proteins involved in molting and cuticle production (e.g., structural constituent of cuticle, collagen-containing ECM, etc.) (**Fig. 3A**). These findings support the notion that normal hypodermal differentiation and function are disrupted or significantly delayed in *eff-1* mutants. Combined with the tissue enrichment analysis (**Fig. 3C**), these data suggest that the changes in gene expression observed in fusion-defective worms occur primarily among genes and pathways associated with the epidermis and epidermal functions relative to other cell and tissue types. To this point, intersecting all shared DEGs with genes with detectable expression in the L1 epidermis found 89.6% of DEGs were present, strengthening the claim that most of genes found to change their expression are present in the larval epidermis.

**Figure 3.**
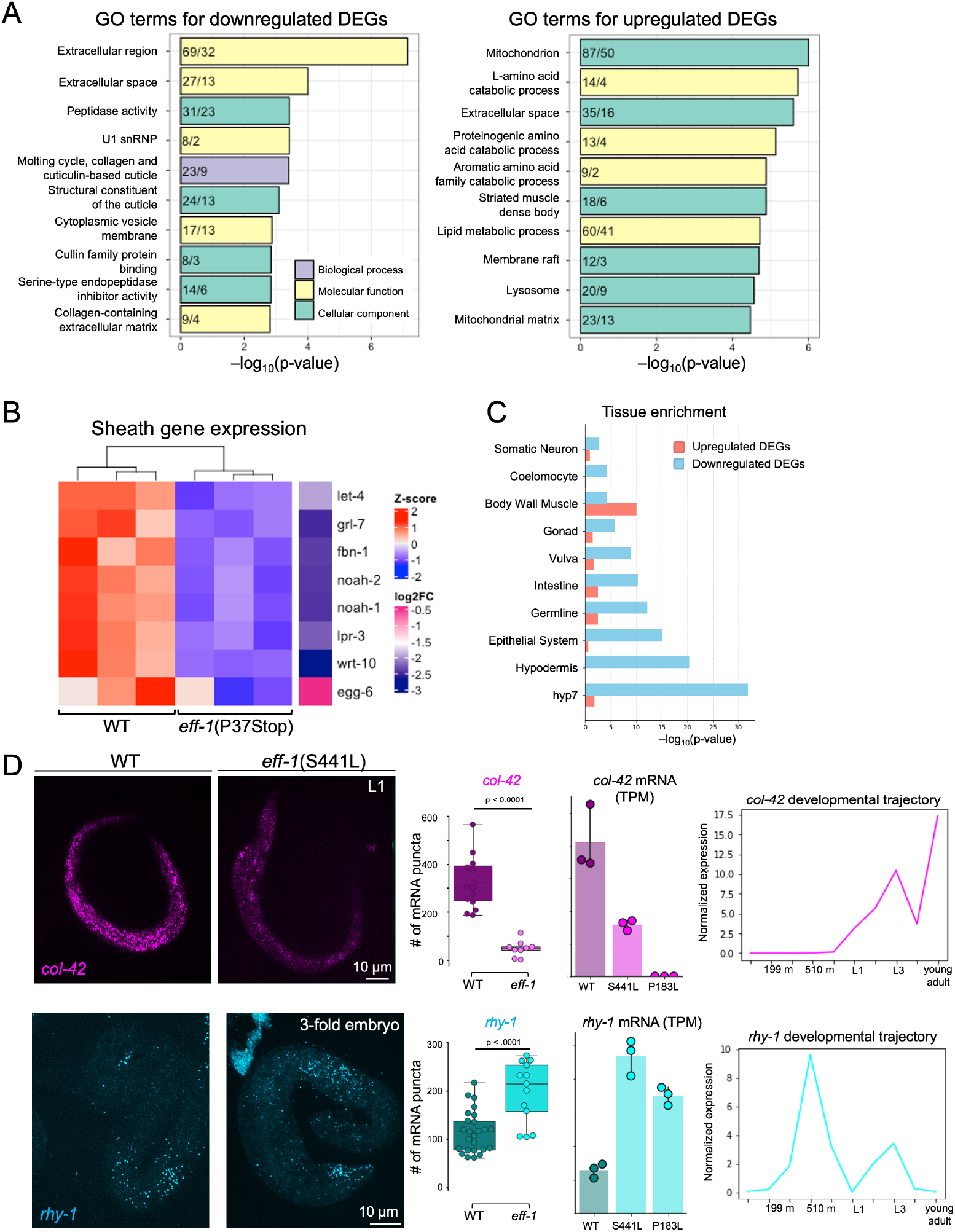
The null *eff-1* mutation affects diverse functions. A) GO term enrichment using topGO with DEGs from *eff-1*(P37Stop) RNAseq. Colors indicate GO term category as shown in the key. Numbers associated with each bar indicate the observed/expected genes for each GO term displayed. B) Heatmap of z-score–transformed mean expression and log_2_(FC) values for selected sheath genes in WT and *eff-*1(P37Stop) L1s. C) Tissue enrichment analysis of DEGs based on gold-standard small-scale enrichment experiments (37). DEGs were intersected with lists of tissue-specific genes, and p-values were calculated based on hypergeometric significance. D) Representative images of *col-42* and *rhy-1* smiFISH hybridization in WT and e*ff-1*(S441L) L1s and 3-fold-stage embryos, respectively. mRNAs were quantified based on the number of fluorescent puncta. RNAseq counts (transcripts per million; TPM) for WT and *eff-1* mutants for *col-42* mRNA at L1 larvae and rhy-1 mRNA in embryos. Developmental trajectory of *col-42* and *rhy-1* expression in WT animals taken from modENCODE expression data (39). Error bars represent standard error; minimum of nine animals per condition.

In addition to the curated set of GO terms on Wormbase, we noted an enrichment in embryonic precuticle-or “sheath”-associated genes that accompanied the decrease in L1-specific genes (**Fig. 3B**). The sheath is the first precuticle formed during embryonic development prior to secretion of the first cuticle (35-38). The presence of precuticle gene expression at the L1 stage in *eff-1* mutants suggests a delay in the transition from precuticle to cuticle gene expression programs. This finding, coupled with the observed downregulation of L1-specific collagen genes, points toward an overall impairment or delay in the transcriptional wiring necessary for proper L1 epidermal maturation and function.

To independently validate and visualize changes in the expression of specific DEGs, we used single-molecule inexpensive fluorescence *in situ* hybridization (smiFISH) of two DEGs identified in our RNAseq data, *col-42* and *rhy-1. col-42* encodes a structural collagen largely restricted to larvae and adults and was significantly downregulated in both *eff-1*(S441L) and *eff-1*(P183L) mutants (**Fig. 3D**). *col-42* is expressed during early larval development, and smiFISH imaging showed almost no detectable signal in *eff-1* mutants at L1 (**Fig. 3D**). Conversely, *rhy-1* is a transmembrane acyltransferase that is expressed in the embryonic epidermis (40, 41). Consistent with the transcriptomic data, the number of *rhy-1* mRNA fluorescent puncta was also elevated in *eff-1* mutants (**Fig. 3D**).

Although large changes in the transcriptomes of *eff-1* mutants were evident, such changes may be due to direct or indirect effects caused by the disruption of epidermal cell functions. To distinguish between these possibilities, we looked for overlap between gold-standard lists of tissue-specific genes from small-scale single-tissue experiments and the *eff-1* DEGs (42). This analysis showed a significant enrichment among downregulated DEGs for transcripts associated with the hyp7 syncytium, hypodermis, and epithelial system, over other non-syncytial tissues (**Fig. 3C**), consistent with a model in which unfused hyp7 progenitors fail to upregulate genes found in the L1 syncytial hypodermis. This suggests that the observed transcriptomic changes are, in large part, the direct consequence of an unfused epidermis rather than of a systemic developmental delay. Collectively, our data support the central finding that fusion plays a critical role in promoting the timely differentiation and function of the epidermis based on hallmark changes in gene expression.

### *eff-1* mutant transcriptional changes are consistent with a developmental delay

In addition to shared pathways and functions associated with the DEGs isolated across *eff-1* mutations, we found a striking enrichment in the DEGs associated with the transition from embryonic to larval development. By examining the developmental trajectory of DEGs at timepoints throughout the WT worm lifecycle from modENCODE (39, 43), we found an enrichment in L1-associated genes in the downregulated DEG set, along with upregulated DEGs showing substantial enrichment for genes normally expressed in the late embryo (**Fig. 4A, D**). Furthermore, genes that were not differentially expressed (FDR > 0.05) in the *eff-1* mutants were predominantly those for which expression levels remain relatively stable across normal development.

**Figure 4.**
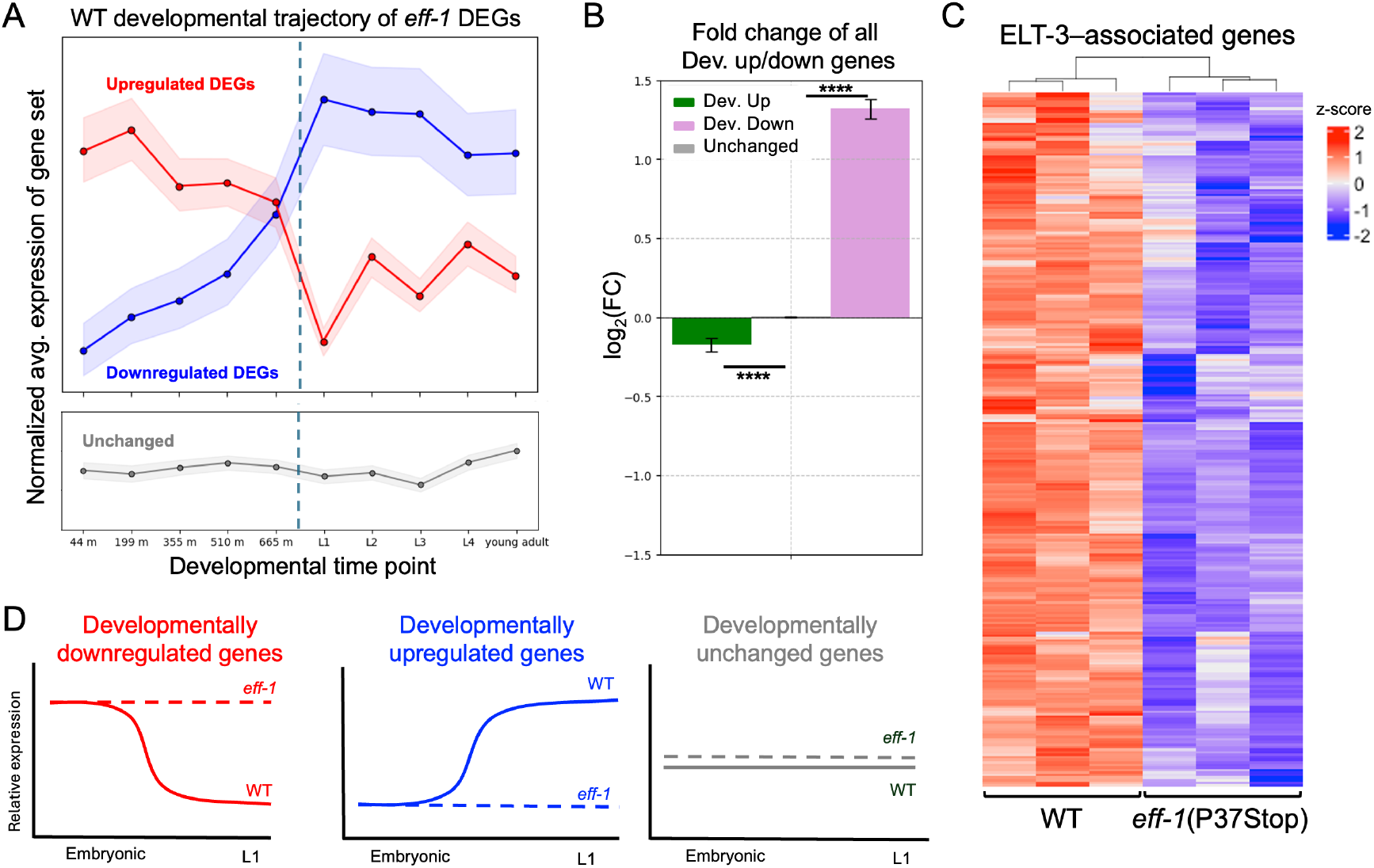
Fusion mutants fail to acquire the L1 epidermal transcriptome. Normal (WT) developmental trajectories of genes that were upregulated (red) or downregulated (blue) in *eff-1* mutants. Shaded region denotes 95% confidence interval. Fusion-defective transcriptomes show a marked increase in embryonic transcripts with a concurrent downshift in larval-associated transcripts. Dashed line indicates an approximate hatching time of 800 min. B) Genes were binned using a linear regression of modENCODE data to determine transcripts that went up (green), down (purple) or stayed consistent (gray) across normal WT development. Average log_2_–transformed fold change data of *eff-1*(P37Stop) RNAseq data for each category of genes are shown. Error bars represent standard error, and p-values calculated using Mann-Whitney U test, ****P < .00001. C) Heatmap of z-score normalized DEG expression for genes identified as ELT-3 regulated from the TF2DNA dataset (49). D) Schematic of WT and *eff-1* mutant gene expression profiles across development, showing a lack of transition from the embryonic to larval transcriptome in fusion defective *eff-1* mutants.

Collectively, these observations strongly suggest that the normal transcriptional shift from late-embryonic to L1 larval gene expression programs is substantially delayed in worms deficient for EFF-1–mediated fusion.

To further investigate the hypothesis that *eff-1* mutant L1s inappropriately maintain high expression of embryonic genes while failing to upregulate larval-specific genes, we performed a linear regression analysis. This analysis used WT expression data across multiple embryonic and larval timepoints to define gene sets that are developmentally upregulated or downregulated during the transition from late-embryonic to larval stages. We then compared the average fold changes of these developmentally dynamic gene sets to *eff-1*(P37Stop) mutants. This comparison revealed a net increase in the expression of developmentally downregulated (i.e., typically embryonically enriched) genes and a corresponding decrease in the expression of developmentally upregulated (i.e., typically larvally enriched) genes, irrespective of whether individual genes met the strict criteria for differential expression (**Fig. 4B**). This broader analysis reinforces the conclusion that the absence of cell fusion leads to a significant delay in the acquisition of the appropriate L1 transcriptome.

The transition from an embryonic to larval transcriptome is facilitated by many transcription factors, including ELT-3, an embryonically enriched GATA transcription factor with roles in the transcriptional activation of epidermis-specific genes and developmental pathways (44-46). Previously, an *elt-3* reporter construct was found to be expressed in only a subset of hypodermal precursor cells, and thus it has been used as a reporter for fusion, or lack thereof, in *eff-1* mutant worms (47). Intriguingly, we found a large swath of ELT-3 regulated genes identified through the TF2DNA database as significantly downregulated in *eff-1* mutant worms (**Fig. 4C**). Examination of the ELT-3 chromatin immunoprecipitation sequencing (ChIPseq) signal across all up, down, and unchanged genes also showed an enrichment of downregulated genes (**Fig. S2**). The observed downregulation of ELT-3 target genes in *eff-1* mutants provides a potential mechanistic link between the failure of cell fusion and the observed developmental delay. Namely, cell fusion may be necessary for the appropriate stage-specific activity of key developmental regulators such as ELT-3, which may be expressed in only a subset of cells contributing to epidermal syncytia.

Taken together, our work underscores the critical role of EFF-1–mediated cell fusion in the developmental progression of *C. elegans* and highlights its importance for the timely activation of the L1 larval transcriptome. Our findings suggest that the process of fusion itself may be a key mechanistic trigger promoting the shift from the embryonic to the L1 transcriptional landscape in fusion-driven syncytia such as hyp7. Although these results highlight the importance of syncytial formation for the proper expression of developmental genes, precisely how fusion orchestrates transcriptional maturation remains an open question. Future investigations will be needed to unravel the molecular mechanisms involved and to identify specific factors that may drive the identity and transcriptional programs of developing syncytial nuclei.

### Materials and Methods

### Strains used and maintenance

*C. elegans* strains were maintained according to standard protocols and were propagated at 22°C. The two *eff-1* mutant strains used here were WH171 (*eff-1*(*oj55*); jcIs1 *[ajm-1::GFP + unc-29(+) + rol-6(su1006)]*) and BP76 (*eff-1*(*hy21*); jcIs1 *[ajm-1::GFP + unc-29(+) + rol-6(su1006)]*). MH1384 (*[kuIs46 (ajm-1::GFP; unc-119(+))]*) was used to compare WT AJM-1::GFP expression to that in e*ff-1* mutants. The P37Stop strain as generated using CRISPR via the method described in Ghanta et. al 2021 (48). Briefly, RNPs loaded with gRNA (GGTGTCTTGGAACAGTGTGG) were injected along with repair template designed to add two stop codons along with an SpeI recognition site into the gonads of animals expressing *kuls46*[AJM-1::gfp] and *bqSi640*[*dpy-7p*::FRT::mCherry::*his-58*::FRT::GFP::*his-58*].

### Developmental timing analysis

Time to hatch was determined by plating young adult worms, allowing them to lay eggs for 2 hours, and then removing the adults and measuring the percentage of unhatched eggs that remained over time. Time to molt was determined by plating synchronized L1 larvae and watching for cessation of pharyngeal pumping to mark the beginning of the L1–L2 molt. Time to adulthood was measured by plating synchronized L1 larvae and noting the presence of embryos in the gonads. All experiments were carried out at 22°C.

### Microscopy

Confocal images for **Fig. 1** were taken using CellSens on an Olympus Spin-SR, Spinning Disc, Super Resolution System using an inverted Olympus IX83 microscope and Yokogawa W1 spinning disc. smiFISH images were taken of worms fixed in methanol/acetone as described (49). Larvae for **Fig. 1** images were prepared in 10 mM levamisole on 3% agarose pads.

### RNA isolation and RNAseq

L1s were synchronized by bleaching gravid adults and allowing embryos to arrest in M9 buffer. L1s were spun down and lysed for RNA collection via the Direct-zol kit from Zymo. RNA quality was confirmed using an Agilent TapeStation with minimum RINs (RNA Integrity Number) of 8.0. mRNA libraries were constructed using TruSeq Library Prep Kits from Illumina and were sequenced on an Illumina Nextseq P3 flow cell.

### Transcriptomic analysis

Paired-end reads were processed as follows. Low-quality reads were filtered using fastp, and their quality was assessed using fastQC (0.12.1). Reads were aligned using STAR and pseudoaligned using Salmon to the *ce11* reference genome, duplicates were removed using picard, and counts were collected using htseq (50-52). Differential expression was determined using DESeq2 (v. 1.42.1), with counts filtered for >1 TPM across all samples. DEGs were defined as genes with a Benjamini-Hochberg adjusted p < 0.05 and log_2_(fold change) > 0.5 or < –0.5, (processed files in **File S1)**. Heatmaps were generated using the seaborn clustermap() function using Euclidian distance to obtain hierarchal linkages and z-score normalization by rows. Intersectional analysis was done by collecting modENCODE developmental expression data, processed in Hutter and Soh 2016 (39, 43). Upregulated, downregulated, and unchanged classifications were determined by performing a linear regression for expression data across developmental time points and collecting genes with a derived slope of >1 (developmentally upregulated) or <–1 (developmentally downregulated). Genes with a derived slope from 1 to –1 were categorized as unchanged. Raw and processed data files can be found through GEO accession: (GSE301850).

### GO term analysis

GO term enrichment was carried out using topGO (v. 2.54.0) on ontologies from WormBase using the “weight01” algorithm, and significance was assessed with a Fisher’s statistic. Genes found to be differentially expressed (FDR < 0.05) in both S441L and P183L were used as the test set, and the background gene set was specified as genes with a base mean TPM of >1 in all three transcriptomes.

### Tissue enrichment

Gold-standard tissue-specific gene sets were obtained from https://worm.princeton.edu/ (42) and compared to DEGs from the *eff-1* mutant RNAseq data. Fischer’s exact test was used to calculate enrichment significance.

### smiFISH

smiFISH was carried out as described (49). Probe sets were designed against *col-42* and *rhy-1* using Oligostan with FLAP-X extensions (53). The FLAP-X secondary probes were labeled with Quasar 670 from Biosearch Technologies. Quantification was done in FIJI by manually drawing designated regions of interest around embryos and L1s of appropriate age and sufficient fixation, thresholding across experimental replicates, and using the “analyze particles” option.

### ELT-3 ChIPseq analysis

ELT-3–associated genes were obtained from the TF2DNA dataset (54) and compared to our experimentally derived DEGs. L1 ELT-3 ChIPseq data from modENCODE was downloaded as read-depth–normalized bigwig files. Heatmaps were generated with *deeptools* (v 3.5.6) (55). Up- and downregulated gene sets were designated from the *eff-1* P37Stop dataset with the same criteria (adjusted p-value < .05, log2 fold change > 0.5 or < -0.5. Unchanged genes were taken as a random subset of 5000 genes with adjusted p-value > .2.

## Acknowledgements

This work was supported by the National Institutes of Health grants GM136236 and GM125091 (to D.S.F.), GM134885 (to D.L.L.). This project was also supported by an Institutional Development Award (IDeA) from the National Institute of General Medical Sciences of the National Institutes of Health under P20GM103432. We thank Amy Fluet for editing this manuscript, the Nishimura lab for guidance and implementation of smiFISH, the Sundaram and Murray labs for insightful conversation and feedback on the results.

## Author contributions

Conceptualization, OHF, DLL, DSF; Investigation, OHF; Writing – Original Draft, OHF; Writing – Review & Editing, OHF, DLL, DSF; Funding Acquisition, DLL, DSF; Supervision, DLL, DSF.

---

**Figure S1.**
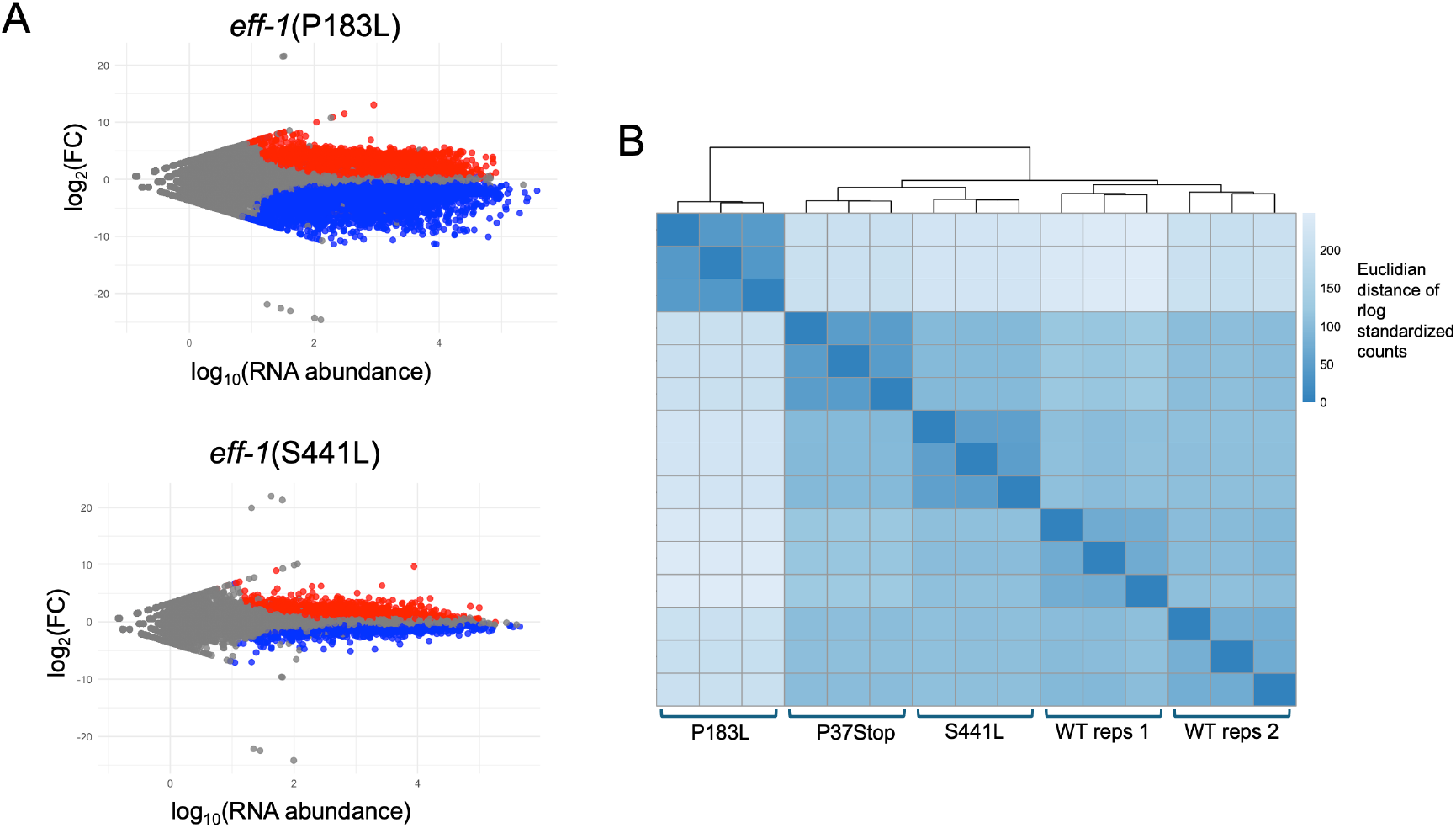
RNAseq of *eff-1* mutants shows substantial overlap in gene expression changes. A) MA plots of *eff-1*(P183L) and *eff-1*(S441L) with DEGs (adjusted p < 0.05) in red for upregulated and blue for downregulated. B) Euclidian distance matrix of regularized log transformed RNAseq counts of *eff-1* mutants and WT controls from multiple sequencing runs.

**Figure S2.**
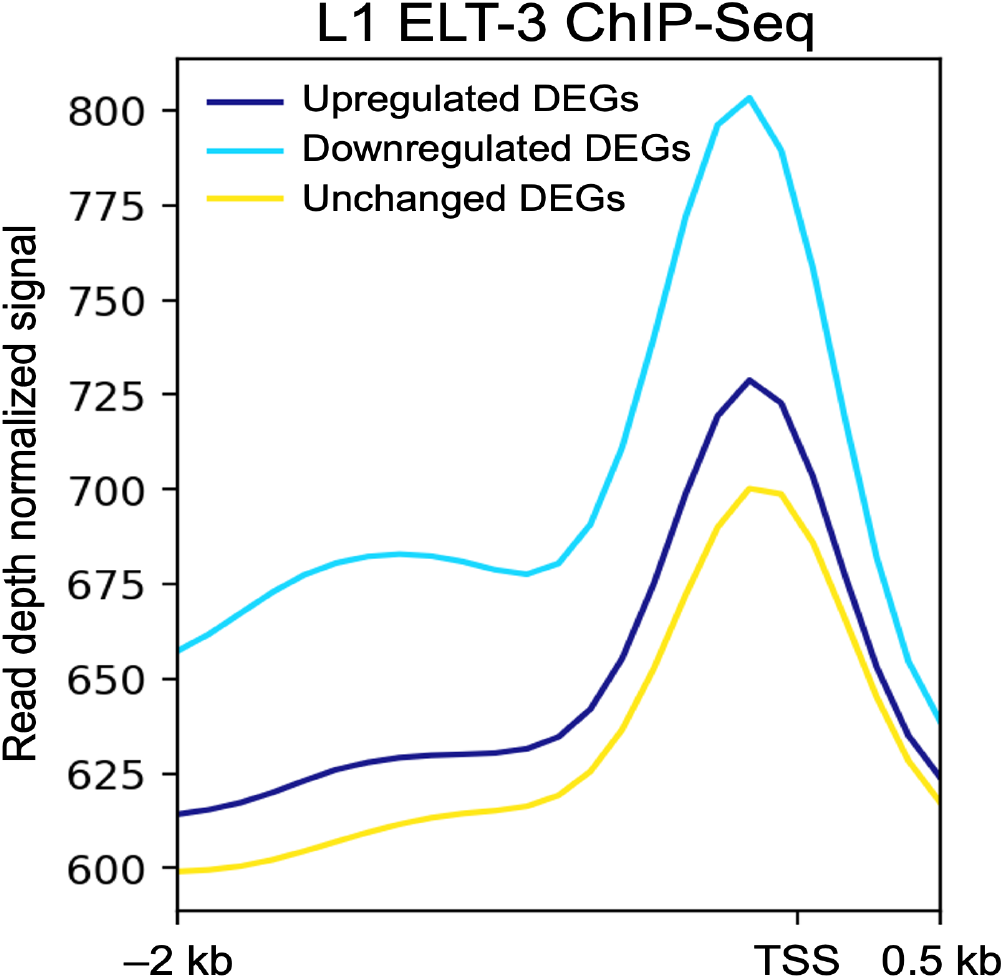
ELT-3 ChIPseq signal across *eff-1* differentially expressed genes. L1 ELT-3 ChIPseq profile across DEGs from *eff-1*(P37Stop). ELT-3 ChIPseq signal was derived from modENCODE data and plotted for a region spanning 2 kb upstream and 500 bp downstream from the transcription start site (TSS) of upregulated, downregulated, and unchanged genes.

## Notes

### Competing Interest Statement

The authors have declared no competing interest.

### Summary of Updates

Additional experiments and analyses were carried out along with edited writing.

