## Supplementary figures and images for "Epidermal cell fusion promotes the transition from an embryonic to a larval transcriptome in *C. elegans*"

### Fig S1

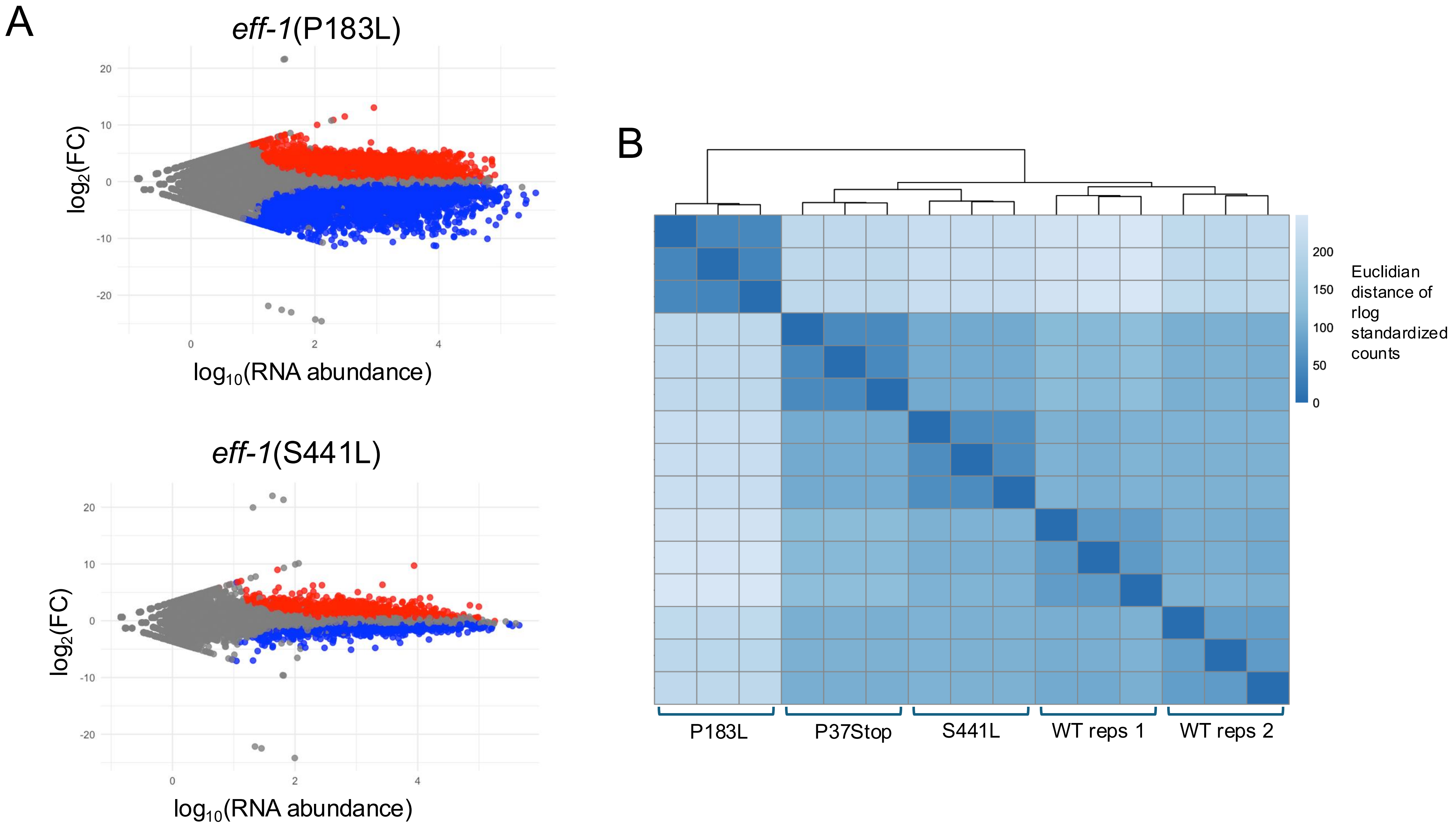

### Fig S2

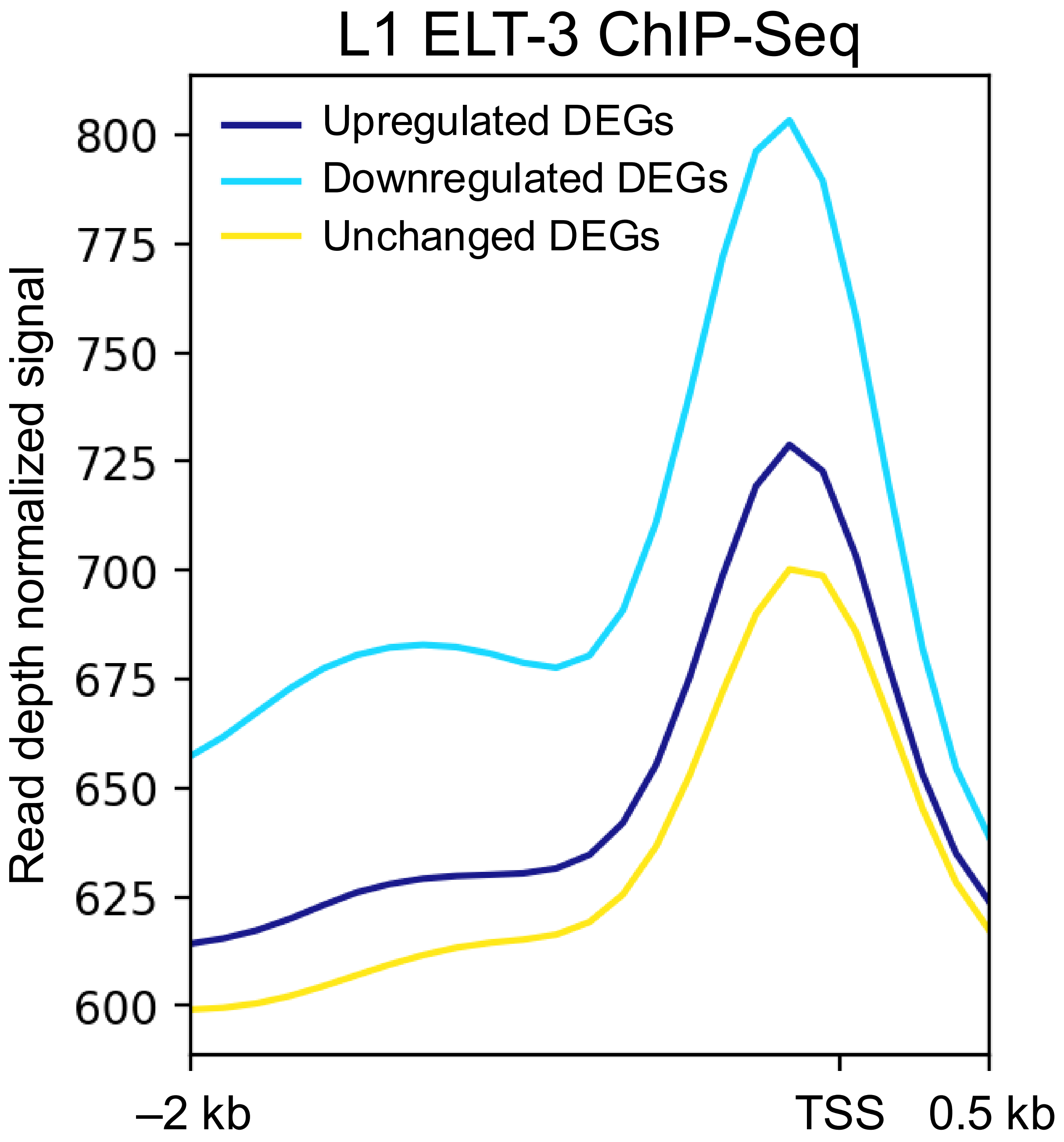
